# Positional frequency chaos game representation for machine learning-based classification of crop lncRNAs

**DOI:** 10.1101/2025.06.03.657533

**Authors:** Athanasios Papastathopoulos-Katsaros, Zhandong Liu

## Abstract

Alignment-based methods are fundamental for sequence comparison but are often computationally prohibitive for large-scale genomic analyses. This limitation has spurred the development of quicker, alignment-free alternatives, such as k-mer analysis, which are crucial for studying long noncoding ribonucleic acids (lncRNAs) in plants. These lncRNAs play critical roles in regulating gene expression at both the epigenetic and transcriptomic levels. However, existing alignment-free approaches typically lose positional information, which can be vital for achieving accurate classification. We propose positional frequency chaos game representation (PFCGR), a novel encoding that improves the traditional frequency chaos game representation (FCGR) by incorporating four statistical moments of k-mer positions: mean, standard deviation, skewness, and kurtosis. This creates a multi-channel image representation of genomic sequences, enabling machine learning models such as Logistic Regression, Random Forests, and Convolutional Neural Networks to classify plant lncRNAs directly from raw genomic sequences. Tested on seven major crop species, our PFCGR-based classifiers achieve classification accuracies comparable to or exceeding those of the computationally intensive DNABERT-based model [1], while requiring 80% to 95% less computational time. These results demonstrate PFCGR’s potential as an efficient and accurate tool for plant lncRNA identification, as well as its ability to facilitate large-scale computational studies in genomics.

## I. Introduction

Alignment-based methods are essential for analyzing biological sequences but require significant computational resources. This need has accelerated the development of faster, alignment-free options, like k-mer analysis, which are vital for the study of different biological organisms. In plant biology, alignment-free methods show great promise for identifying long non-coding RNAs (lncRNAs) in crop species, offering significant advantages over traditional alignment-based approaches.

LncRNAs, defined as RNA molecules exceeding 200 nucleotides in length, are a critical component of the plant transcriptome with no coding potential but significant regulatory functions [2]–[4]. They are transcribed by RNA Polymerase II and are often polyadenylated, though significant exceptions exist. Plant genomes, unlike those of yeast and mammals, also utilize the plant-specific RNA polymerases Pol IV and Pol V to produce distinct lncRNAs, some of which are non-polyadenylated. These include upstream noncoding transcripts (UNTs) in *Arabidopsis* that are analogous to mammalian promoter upstream transcripts (PROMPTs), suggesting a nuanced regulatory landscape [5].

LncRNAs are involved in a myriad of regulatory roles across various stages of plant development and are differentially expressed across tissues, reflecting their specialized functions. Notably, lncRNAs influence key developmental processes, such as the timing of flowering [6], [7], and are crucial for plants’ resilience to environmental stresses [8]–[12].

In terms of functionality, lncRNAs modulate gene expression through a variety of mechanisms, including epigenetic modifications [13]–[15]. For instance, in rice, the lncRNA LAIR notably increases grain yield by upregulating the LRK gene cluster, which leads to larger and more numerous panicles [16]. Another lncRNA, Ef-cd, sourced from the antisense strand of the OsSOC1 locus, significantly shortens the maturity period for late-maturing rice varieties without reducing yield, showcasing lncRNAs’ ability to fine-tune plant growth under cultivation constraints [17]. These advancements in lncRNA research are paving the way for significant contributions to agricultural biotechnology, including the development of genetically engineered crops with increased yields, improved resistance to pests and diseases, and greater environmental stress tolerance. Thus, understanding and predicting lncRNAs remain crucial in modern plant science and biotechnology. For additional implications of lncRNAs in physiological processes, we refer the reader to [18].

To address the challenges of existing methods, we introduce a novel approach for classifying plant lncRNAs based on genomic sequences, using machine-learning techniques such as Logistic Regression (LR), Random Forest (RF), and Convolutional Neural Networks (CNN). Our approach uses Frequency Chaos Game Representation (FCGR) and our innovative improved version, “positional FCGR” (PFCGR), which incorporates the following statistical features of the positions of all the k-mers of the sequences: mean, standard deviation, skewness, and kurtosis. Therefore, the aims of this study are: (i) to address FCGR’s limitation of losing positional information; (ii) to develop and evaluate PFCGR-based machine-learning models for the identification of plant lncRNAs directly from genomic sequences; (iii) to compare the accuracy and computational efficiency of PFCGR-based models against the very accurate and much more complex DNABERT-based method across seven crop species.

## II. Related work

Bioinformatics pipelines for lncRNA prediction often combine custom scripts and tools like CPC2 [19], LncMachine [20], CPAT [21], PlncPRO [22], PLEK [23] and PLEKv2 to filter and estimate the coding potential of transcripts from transcriptome datasets. These methods typically involve manually defining features from transcripts, which are then used in machine-learning models to classify transcripts as mRNA or lncRNA [25], [26]. However, the reliance on handcrafted features can introduce bias and affect classification accuracy [27], [28], while the training of models depends heavily on transcriptome data from sources like CANTATAdb [29], PLncDB [2], GreeNC [30], [31] and AlnC [32]. For this reason, a shift towards models that utilize genomic references could improve the detection of lncRNAs, particularly given their tissue- or stage-specific low expression rates.

One such work [1] is based on DNABERT [33], a specialized adaptation of the language BERT model [34] tailored for DNA sequences. It addresses the binary classification task of distinguishing true lncRNA from synthetic lncRNA sequences, as explained in more detail in subsequent sections. DNABERT is capable of performing sequence classification by encoding DNA sequences into k-mer tokens, which allows it to capture intricate biological features. This is achieved through the transformer architecture and the attention mechanism [35], which unfortunately comes with a significant computational cost and the black-box nature of the model. While other recent transformer-based approaches like LncRNA-BERT [36] and LoRA-BERT [37] have demonstrated strong performance for lncRNA classification, they present similar computational challenges and are primarily trained on human data with limited evaluation on plant species. For these reasons, we only compare our method against [1] to ensure a fair and equal comparison on the exact same dataset. For a more extensive discussion on related lncRNA work, refer to [38] and [1].

Lastly, we acknowledge that the loss of positional information, the main limitation of FCGR, has been a subject of previous research. Methods have been developed to decode positional information from FCGR images using techniques such as wavelet transformation (Frequency Chaos Game Signal [39]) or by analyzing correlations among k-mers [40]. However, these approaches differ significantly from ours, as none have implemented a similar strategy of using statistical features of k-mer positions to capture and utilize this information. By introducing PFCGR, we propose a distinct and novel way to address this long-standing limitation, and we choose to benchmark it to the challenging task of crop lncRNA classification.

## III. Methodology

### A. FCGR encoding

The Chaos Game Representation (CGR), originally used to construct fractals from random inputs, was later adapted to encode DNA sequences [41]. For readers unfamiliar with CGR, a simple illustration is provided in Figure 1. In this work, we use the Frequency CGR (FCGR), a discrete matrix equivalent of CGR that counts k-mer frequencies at each position. Unlike traditional encoding methods such as one-hot encoding, FCGR encodes nucleotide sequences into images/numerical data, standardizing sequences of varying lengths into images of the same size. For further reading, see [42].

**Fig. 1.**
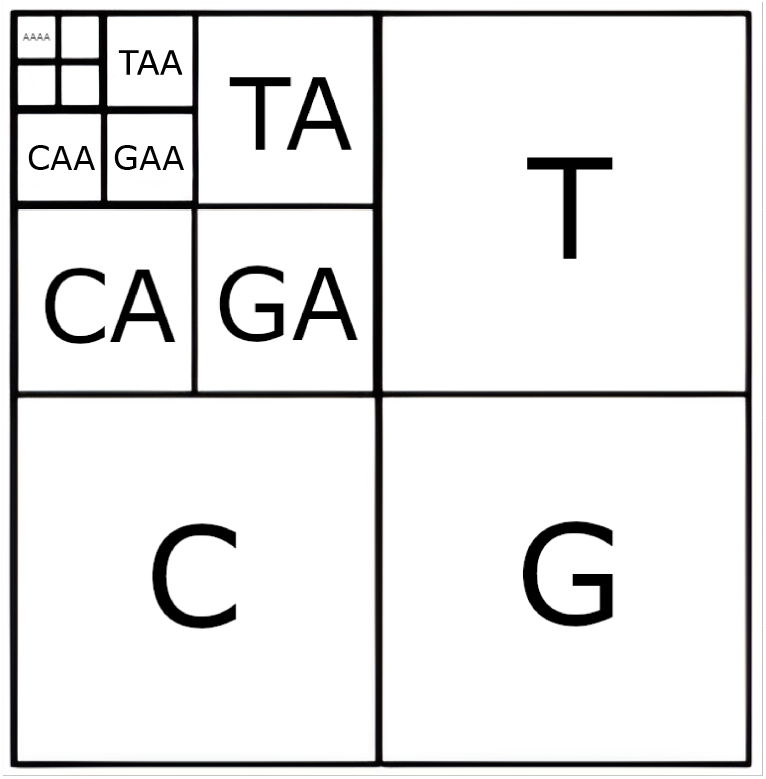
Illustration of the CGR algorithm: The square is initially divided into four equal parts, each representing one of the four nucleotides found in DNA: Adenine (A), Thymine (T), Cytosine (C), and Guanine (G). Each quadrant is assigned to one nucleotide. Each of these initial quadrants is further subdivided into four smaller squares. This division process respects the nucleotide of the larger square from which it is derived. For example, the square for Adenine (A) is subdivided into AA, TA, CA, and GA. This method applies recursively to all quadrants. With each iteration, the squares become increasingly specific to particular sequences of nucleotides. For instance, the top left quadrant, after the second iteration, shows further subdivisions like AAA, TAA, CAA, and GAA, each representing a specific 3-mer, and the grid in the top-left corner of this figure represents the 4-mer AAAA. Finally, to map a DNA sequence onto this square, we begin at the center and move to each subsequent quadrant based on the next nucleotide in the sequence. The position within the square becomes more precise with each nucleotide added, tracing a path through increasingly smaller quadrants.

In a nutshell, the idea behind FCGR is to map k-mers onto a unit square by recursively dividing the square into smaller regions according to the sequence of bases in each k-mer. Each base (A, C, G, T) determines which quadrant of the current region to select, and this subdivision continues for all positions in the k-mer, resulting in a unique location within a 2^*k*^ *×* 2^*k*^ grid corresponding to that k-mer. The FCGR image is constructed by plotting each k-mer’s frequency on that grid, which is normalized to compute pixel intensities. In this study, k ranges from 3 to 6. Figure 2 shows FCGR images from the *Zea Mays* dataset, highlighting some key regions that could potentially aid in sequence classification.

**Fig. 2.**
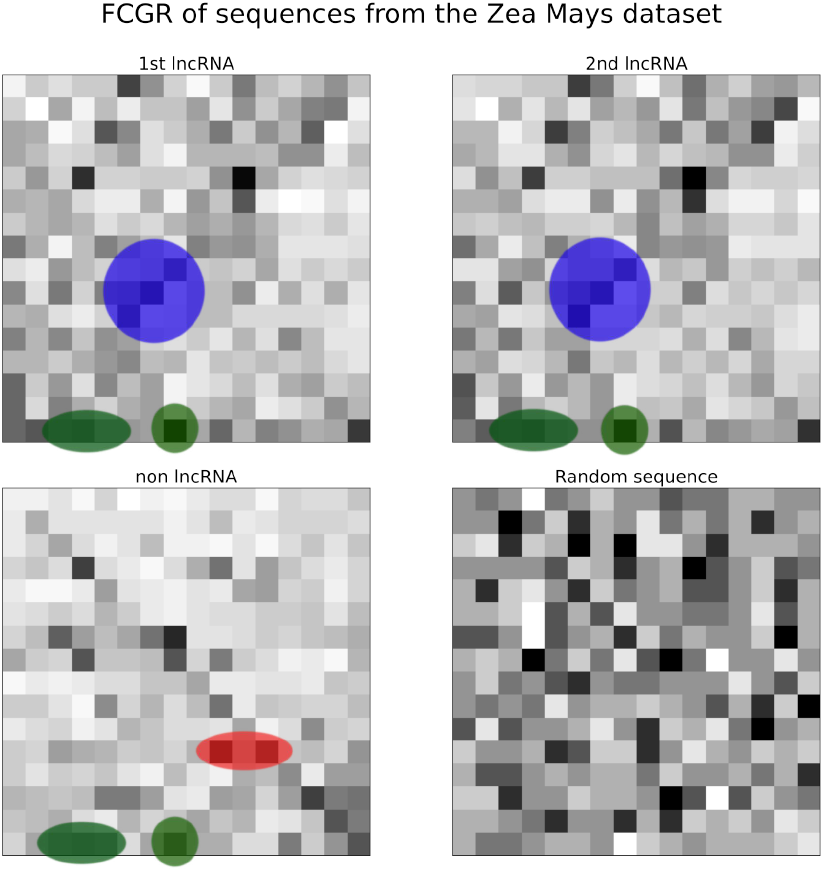
FCGR images with k=4 are displayed for four sequences: two lncRNAs, one non-lncRNA, and one random sequence. Green circles highlight patterns common to both lncRNAs and the non-lncRNA, indicating shared genomic motifs. Blue circles indicate patterns unique to lncRNAs, suggesting motifs specific to lncRNA. Red circles highlight patterns exclusive to the non-lncRNA sequence. Not all patterns are marked to maintain visual clarity in the images.

FCGR has shown success in various applications, such as detecting COVID-19 [43], [44], distinguishing human coding and non-coding regions [45], and predicting antimicrobial resistance [46]. It has also been shown to outperform traditional one-hot encoding in bacterial 16S rRNA classification [47]. Recently, a new FCGR-based representation was developed based on ranked frequencies instead of raw, which was integrated with Transformers and showed improved performance, particularly in low-coverage genomic data [48]. Lastly, reference [49] applied FCGR in unsupervised clustering of DNA sequences using CNN combined with twin contrastive learning, demonstrating high clustering accuracy across viral genomes and other genomic datasets.

### B. Positional FCGR encoding

One major drawback of FCGR is that it does not retain positional information. For this reason, we extend the FCGR method by adding positional information (PFCGR) through statistical measures of the k-mer positions, including mean, standard deviation, skewness, and kurtosis. These are computed using Welford’s online algorithm [50], reducing memory requirements and computational overhead. Each statistical measure generates a separate 2^*k*^ *×* 2^*k*^ image or is kept as tabular data, as explained below. The intensity of each pixel corresponds to the statistical value of the respective k-mer and is normalized by min-max normalization. This added positional context improves FCGR by capturing more complex patterns in genomic sequences, where k-mer placement matters.

There are two ways to encode these added statistical measures. On the one hand, we can organize the extracted features into tabular data. For FCGR, each column represents a k-mer (e.g., “AAT”), and each cell contains its frequency in each sequence of the dataset. For PFCGR, the columns are labeled with both the k-mer and the statistical descriptors (Frequency, Mean, Standard Deviation, Skewness, Kurtosis). The data in this format can then be inputted into LR and RF models. On the other hand, the data in image format (which can be thought of as either five images or one image with five channels) can be inputted into CNN models, which are effective in identifying patterns in the generated images, even across channels. For missing k-mers, we assign zero frequencies in the FCGR matrix. We experimented with different imputation strategies for handling the rest of the missing statistics in the PFCGR matrices, including filling with zeros, using -1 as a sentinel value, and adding an additional binary masking channel indicating presence or absence of a k-mer. Since all approaches yielded similar performance, we opted to use zeros for simplicity. This issue does not affect the LR and RF methods, as the tabular format does not contain those columns.

### C. Model architectures and training

In selecting our classifiers, we prioritized simplicity and computational efficiency to establish a proof of concept for PFCGR’s effectiveness, rather than pursuing complex architectures for absolute maximum performance. This study represents the first application of PFCGR, and specifically using PFCGR for lncRNA classification, making it essential to demonstrate the method’s viability using straightforward models. Therefore, LR and RF were chosen for their low computational cost and ease of computing feature importance, while CNNs were included for their ability to capture complex patterns in image-based data. Our CNN architecture for FCGR includes, in this order, a convolutional layer, followed by a pooling layer, a fully-connected hidden layer, and a final sigmoid output neuron. All other layers use ReLU activation functions. The difference for PFCGR is that the five images are processed separately by one convolutional layer before merging their outputs to the rest of the neural network, which has the same architecture as the single-image FCGR setup. This is intended to capture unique features of each image before further processing.

Each species has a dedicated classifier, individually trained in species-specific datasets, to accurately distinguish true lncRNA from synthetic non-lncRNA sequences. For LR, we applied L2 regularization, which demonstrated more robust performance compared to LASSO. For RF, we optimized the number of trees and their maximum depths using grid search. Specifically, the number of trees was varied between 250 and 1250, while the maximum depth ranged from unrestricted to a limit ranging from 10 to 40 to reduce overfitting. Lastly, for the CNN, the number of filters and filter sizes in the convolutional layers were optimized via grid search. The max pooling layer’s window size and stride were also tuned, along with the number of neurons in the fully connected layer. Specifically, the number of filters was varied between 16 and 128, while kernel sizes ranged from 1×1 to 8×8. Pooling window sizes and strides were explored from 2×2 up to 6×6 to identify the optimal configuration. The model was trained using binary cross-entropy loss with the Adam optimizer (learning rate = 0.001), batch sizes ranging from 8 to 32, and for 7 to 10 epochs. To prevent overfitting and ensure reliable performance, stratified 5-fold cross-validation was applied to all models. The CNN models were trained on an NVIDIA A30 GPU to reduce training time.

### D. Computational complexity of PFCGR compared to DNABERT

To compare the computational complexity of PFCGR to DNABERT’s, we consider both feature extraction and classifier complexity, using RF for illustration. For a sequence of length *N* and a fixed k-mer size, feature extraction involves iterating over the sequence once (sliding a window of length *k*), with each *k*-mer requiring *𝒪* (1) time because of hashing and Welford’s online algorithm ability to compute the statistical measures on-the-fly. Thus, the total complexity for generating the PFCGR feature vector is *𝒪* (*N*). For *M* training sequences, each with a feature vector of length *≈* 5 *·* 4^*k*^, the RF complexity with *T* trees and tree depth d_tree_ is *𝒪* (*M ×* (5*·*4^*k*^) *×T ×* d_tree_). The term 4^*k*^ represents the maximum possible number of unique k-mers. For small *k*, the feature vector size is manageable. For large *k*, the number of unique k-mers present is typically much smaller than 4^*k*^. In this case, the number of unique k-mers is bounded by the sequence length and therefore, while the theoretical upper bound is exponential in k, the practical complexity remains tractable for both small and large values of k. In contrast, DNABERT, has a computational complexity of 𝒪 (*L ×* (*N* ^2^ *×d* + *N× d*^2^)) for a sequence of length *N*, where *d* is the embedding size and *L* is the number of feedforward layers. The self-attention mechanism introduces a quadratic complexity on *N*, which makes it significantly more computationally demanding than PFCGR. This is particularly problematic for the long sequences in our study (median length 1500-2200 nucleotides, see Table I), as DNABERT’s performance and efficiency degrade with increasing sequence length due to the high computational cost and challenges in capturing long-range dependencies.

**TABLE I.** Median sequence lengths and label counts for each crop species.

| Species | Median seq. length | lncRNA labels | Non-lncRNA labels |
| --- | --- | --- | --- |
| A. thaliana | 1,514 | 3,467 | 3,464 |
| B. napus | 1,854 | 8,544 | 7,442 |
| B. oleracea | 1,832 | 5,350 | 4,997 |
| B. rapa | 2,070 | 7,075 | 6,882 |
| G. max | 2,143 | 2,320 | 2,288 |
| O. sativa | 1,943 | 2,335 | 2,334 |
| Z. mays | 1,978 | 8,889 | 8,747 |

A direct comparison of training times between PFCGR-based RF and CNN models relative to DNABERT is provided in Table II. Our approaches are about 80% to 95% faster, with the exact speedup varying by classifier and dataset. It is important to acknowledge that this comparison is complex and our PFCGR implementation has room for further optimization. DNABERT is designed to exploit high-performance hardware, taking advantage of large-scale GPUs, mixed-precision training, and optimized matrix operations. In contrast, our current PFCGR implementation does not fully capitalize on these advanced GPU optimizations. These discrepancies in hardware and software optimizations can make a direct, apples-to-apples comparison of raw execution times challenging. Even with these factors making DNABERT’s times appear closer to our current implementation than they are in reality, our results still convincingly demonstrate the superior computational efficiency of the PFCGR-based methods.

**TABLE II.** Comparison of the training times(in minutes) fordnabert, rf withpfcgr, andcnn withpfcgr. all models were trained on the complete dataset using their respective optimal parameters(fordnabert we refer the reader to[1]) using*k* = 6. dnabert andcnn models utilized annvidia a30 gpu, while therf model was trained on anintelxeonsilver4314 cpu. cross-validation and inference times are excluded.

| Species | RF w/ PFCGR | CNN w/ PFCGR | DNABERT |
| --- | --- | --- | --- |
| A. thaliana | 13 | 3 | 54 |
| B. napus | 34 | 9 | 180 |
| B. oleracea | 24 | 6 | 120 |
| B. rapa | 38 | 9 | 216 |
| G. max | 24 | 8 | 96 |
| O. sativa | 20 | 6 | 84 |
| Z. mays | 45 | 13 | 288 |

### E. Datasets and experimental setup

We used the same data set as described by [1]. In brief, the lncRNA datasets for all crop species (*Arabidopsis thaliana, Brassica napus, Brassica oleracea, Brassica rapa, Glycine max, Oryza sativa*, and *Zea mays*) were sourced from CantataDB 2.0 [29] and PLncDB 2.0 [2]. To enrich the dataset, flanking regions were added both upstream and downstream of each lncRNA sequence to prevent the classifier from merely identifying broad genomic features. A control dataset was constructed by extracting random genomic stretches, explicitly excluding the lncRNA sequences and their flanking regions. This approach increases the complexity of the classification task compared to the simpler mRNA versus lncRNA, encouraging the model to learn discriminative features specific to lncRNA regions within their genomic context. For a comprehensive explanation of the dataset construction, we direct the reader to [1].

In our experiments, we performed binary classification between lncRNA and non-lncRNA sequences separately for each species. This choice was motivated by intrinsic differences in the datasets, such as variations in sequence length distributions, nucleotide composition, and class imbalances. Training separate classifiers per species allows them to better capture features unique to each genome. Table I summarizes the median length of the sequences for each species, showing the scale of the datasets, as well as the class distribution for each dataset. The class distribution varies between species, reflecting differences in available annotated sequences and the size of the control sets. For this reason, to robustly evaluate model performance, we employed stratified 5-fold cross-validation. Each fold was used once as the test set, while the remaining four folds served as training data. This approach ensures that the evaluation is reliable and that each fold contains representative samples of both classes.

#### Metrics

As in [1], model effectiveness was measured using accuracy, precision, recall, and F1 score. True positives, false negatives, and false positives represent correctly identified lncRNAs, misclassified lncRNAs, and falsely identified lncRNAs, respectively.

## IV. Results and discussion

We conducted two sets of experiments to evaluate our method. In the first set, we used LR, RF, and CNN for binary classification between non-lncRNA and lncRNA sequences in the seven species. We applied FCGR and PFCGR and optimized the k-mer size for each species, ranging from 3mers to 6-mers, to evaluate the impact of the additional statistical features and their importance. This set also included a comparison with the DNABERT model [1]. In the second set of experiments, we aimed to improve classification accuracy by applying RF and CNN across all k-mer sizes combined (k=3 to k=6), to further improve the model’s performance and provide a better comparison with DNABERT. This set of experiments served to support the findings of the first.

### A. Performance evaluation using the optimal k-mers

To evaluate whether PFCGR boosts performance over the traditional FCGR method, we apply LR, RF, and CNN, using various k-mer sizes to determine the best accuracy. The range of k-mer lengths considered (k ranging from 3 to 6) reflects a common practice in k-mer-based approaches: smaller k values tend to be overly frequent and less informative, while k values greater than 6 become increasingly rare, resulting in sparse and potentially noisy features that can skew model performance. Thus, the selected range is more balanced.

Before presenting our results, we first want to address any skepticism regarding the utility of PFCGR over regular FCGR. To do that, we present an example from the *Zea Mays* dataset in Figure 3. As it can be seen, while lncRNA and non-lncRNA sequences exhibit similar FCGR patterns, incorporating additional statistical measures reveals significant differences. The four new metrics clearly distinguish between the two sequence types, suggesting that the PFCGR method offers more information, potentially improving lncRNA classification accuracy.

**Fig. 3.**
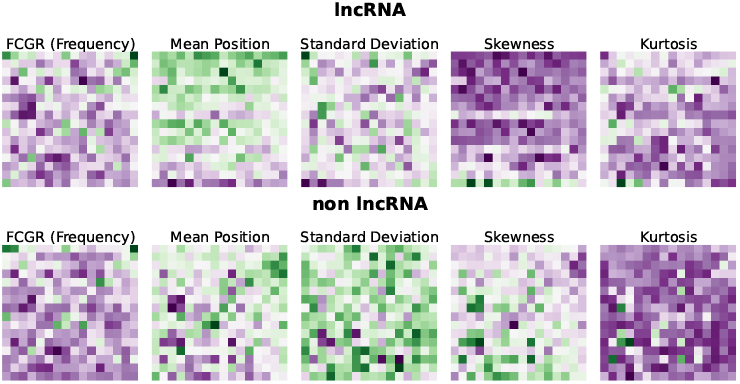
One lncRNA and one non-lncRNA sequence are shown using the standard FCGR and the positional FCGR. The top row displays the lncRNA, while the bottom row depicts the non-lncRNA. Despite very similar FCGRs for both sequences, PFCGRs reveal differences, emphasizing their discriminative power. The color scale ranges from dark purple (indicating low values) to dark green (indicating high values), with all values normalized across the statistics using min-max normalization to ensure consistent comparison.

In Table III, we summarize our results obtained with the optimal k-mer lengths for each species, where “optimal” refers to the k-mer size within the range 3 to 6 that yields the highest classification accuracy for that species. Specifically, a k-mer of length 3 works best for *A. thaliana, B. napus, B. oleracea, rapa*, and *O. sativa*, while lengths of 4 and 6 are optimal for *G. max* and *Z. mays* respectively. All results reported are the averages obtained from stratified 5-fold cross-validation to ensure robustness of the models. Our results demonstrate that PFCGR improves the classification performance for nearly all species compared to FCGR. For certain ones, RF and CNN models achieve results comparable to or better than DNABERT, whose results were fine-tuned separately for each species to optimize performance. In particular, the higher recall across the board suggests that our models are better at identifying lncRNAs, which is critical in most applications.

**TABLE III.** Evaluation metrics for species-specificlr, rf, andcnn models usingfcgr andpfcgr, and a comparison withdnabert, for the binary classification of lncrnas across the seven crop species. the table reports the meanaccuracy, precision, recall, andf1 score from stratified5-fold cross-validation, using the optimal k-mer size for each method.

| Species | Method | Acc. | Prec. | Rec. | F1 |
| --- | --- | --- | --- | --- | --- |
| **A. thaliana** | LR - FCGR | 0.56 | 0.56 | 0.55 | 0.56 |
|  | LR - PFCGR | 0.59 | 0.59 | 0.60 | 0.60 |
|  | RF - FCGR | 0.61 | 0.61 | 0.64 | 0.62 |
|  | RF - PFCGR | 0.65 | **0.67** | 0.60 | 0.63 |
|  | CNN - FCGR | 0.62 | 0.61 | **0.68** | 0.64 |
|  | CNN - PFCGR | **0.66** | 0.66 | 0.65 | **0.65** |
|  | DNABERT | 0.65 | 0.65 | 0.65 | **0.65** |
| **B. napus** | LR - FCGR | 0.55 | 0.54 | 1.00 | 0.70 |
|  | LR - PFCGR | 0.64 | 0.66 | 0.68 | 0.67 |
|  | RF - FCGR | 0.66 | 0.64 | 0.81 | 0.72 |
|  | RF - PFCGR | 0.70 | 0.69 | 0.80 | 0.74 |
|  | CNN - FCGR | 0.66 | 0.64 | **0.81** | 0.72 |
|  | CNN - PFCGR | 0.70 | 0.69 | 0.80 | **0.74** |
|  | DNABERT | **0.75** | **0.75** | 0.74 | **0.74** |
| **B. oleracea** | LR - FCGR | 0.59 | 0.56 | 0.94 | 0.70 |
|  | LR - PFCGR | 0.63 | 0.63 | 0.66 | 0.65 |
|  | RF - FCGR | 0.67 | 0.64 | 0.78 | 0.71 |
|  | RF - PFCGR | 0.71 | 0.71 | 0.74 | 0.73 |
|  | CNN - FCGR | 0.69 | 0.68 | 0.76 | 0.72 |
|  | CNN - PFCGR | 0.68 | 0.65 | **0.85** | **0.74** |
|  | DNABERT | **0.74** | **0.75** | 0.74 | **0.74** |
| **B. rapa** | LR - FCGR | 0.55 | 0.55 | 0.64 | 0.59 |
|  | LR - PFCGR | 0.55 | 0.55 | 0.55 | 0.55 |
|  | RF - FCGR | 0.58 | 0.58 | 0.65 | 0.61 |
|  | RF - PFCGR | **0.60** | **0.60** | 0.67 | 0.63 |
|  | CNN - FCGR | 0.59 | 0.59 | 0.66 | 0.62 |
|  | CNN - PFCGR | 0.56 | 0.55 | **0.80** | **0.65** |
|  | DNABERT | 0.58 | 0.58 | 0.58 | 0.58 |
| **G. max** | LR - FCGR | 0.50 | 0.50 | 1.00 | 0.67 |
|  | LR - PFCGR | 0.62 | 0.63 | 0.63 | 0.63 |
|  | RF - FCGR | 0.70 | 0.68 | 0.77 | 0.72 |
|  | RF - PFCGR | 0.72 | 0.70 | 0.78 | **0.74** |
|  | CNN - FCGR | 0.68 | 0.64 | 0.83 | 0.72 |
|  | CNN - PFCGR | 0.70 | 0.66 | **0.84** | **0.74** |
|  | DNABERT | **0.73** | **0.73** | 0.73 | 0.73 |
| **O. sativa** | LR - FCGR | 0.54 | 0.55 | 0.48 | 0.51 |
|  | LR - PFCGR | 0.57 | 0.57 | 0.57 | 0.57 |
|  | RF - FCGR | 0.61 | 0.61 | 0.61 | 0.61 |
|  | RF - PFCGR | **0.63** | **0.64** | 0.60 | 0.62 |
|  | CNN - FCGR | 0.62 | 0.62 | 0.65 | **0.63** |
|  | CNN - PFCGR | 0.57 | 0.56 | **0.67** | 0.61 |
|  | DNABERT | 0.62 | 0.62 | 0.62 | 0.62 |
| **Z. mays** | LR - FCGR | 0.72 | 0.69 | 0.80 | 0.74 |
|  | LR - PFCGR | 0.72 | 0.72 | 0.74 | 0.73 |
|  | RF - FCGR | 0.79 | 0.75 | **0.88** | 0.81 |
|  | RF - PFCGR | 0.79 | 0.75 | **0.88** | 0.81 |
|  | CNN - FCGR | 0.80 | 0.77 | 0.86 | 0.81 |
|  | CNN - PFCGR | 0.81 | 0.78 | **0.88** | **0.83** |
|  | DNABERT | **0.83** | **0.84** | 0.84 | **0.83** |

Another important observation is that despite logistic regression’s tendency to predict an excessive number of lncRNA cases, which inflates recall, PFCGR moderates these predictions leading to more balanced performance metrics. We also notice some disparities between the CNN and RF models when it comes to the accuracy and F1 score metrics, particularly evident in the PFCGR scenario. In slightly imbalanced datasets like ours, there is no clear answer as to which metric we should use, therefore both the CNN and RF models remain accurate, indicating that for most of the cases PFCGR improves the standard FCGR by providing a more robust classification method for lncRNAs without significantly increasing the computational cost. However, we do realize that in a few cases, PFCGR may actually degrade the performance of the CNN model compared to FCGR, which we attribute to potential overfitting.

Before we move on to the next section, we want to mention a few more observations. The difference in performance between different k-mer lengths is smaller for PFCGR, as shown in Table IV for the example of *A. thaliana*. This suggests that PFCGR adds useful features that contribute to a more robust model. These features refer to the coordinates of FCGR/PFCGR in tabular format. Moreover, feature importance analysis revealed that, for both LR and RF models using PFCGR, the majority of the top 20 significant features were standard deviations and frequencies. This emphasizes the value of incorporating measures of k-mer dispersion along the sequence. To calculate feature importance, we used model-specific criteria: for the LR model, the absolute values of the regression coefficients served as a proxy, while for the RF model, we relied on the average decrease in Gini impurity to measure each individual contribution. Although a few skewness and kurtosis-related features were identified, they were fewer in number, aligning with expectations since they capture more specific distributional aspects and ought to be related to secondary effects. While feature importance analysis is also feasible for CNNs, it tends to be more elaborate and we did not see the need to pursue it here, as we expect the results to be similar to those obtained with LR and RF models.

**TABLE IV.**
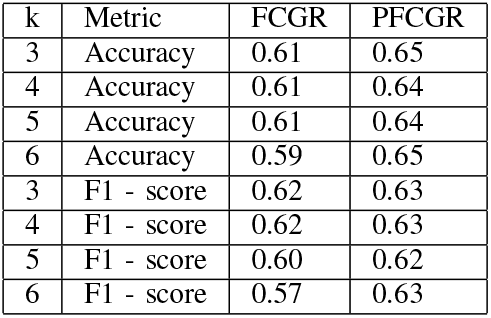
A comparison of the robustness offcgr andpfcgr for different k-mer lengths based on accuracy andf1 score metrics. all results are computed usingrf and the*a. thaliana* dataset and represent the mean performance obtained from stratified5-fold cross-validation.

Lastly, the features showed low correlations (Table V), indicating that they provide complementary information. To compute feature correlations, we calculated the absolute mean correlation and its standard deviation, using Pearson’s correlation coefficient as a measure. This low correlation among features is critical, as it reduces the likelihood of multicollinearity, which can skew the model performance.

**TABLE V.** Mean absolute correlations(mean abs cor.) and their standard deviations(st.d. of the cor.) among all features fordifferent k. all results computed usingrf and the*a. thaliana* dataset.

| k | Statistic | FCGR | PFCGR |
| --- | --- | --- | --- |
| 3 | Mean abs cor. | 0.20 | 0.06 |
| 3 | St.d. of the cor. | 0.15 | 0.08 |
| 4 | Mean abs cor. | 0.10 | 0.03 |
| 4 | St.d. of the cor. | 0.10 | 0.05 |
| 5 | Mean abs cor. | 0.05 | 0.04 |
| 5 | St.d. of the cor. | 0.06 | 0.05 |
| 6 | Mean abs cor. | 0.03 | 0.06 |
| 6 | St.d. of the cor. | 0.04 | 0.06 |

### B. Performance evaluation using all k-mers combined

In our earlier experiments, we observed that the performance did not improve consistently with increasing k-mer size, highlighting the complex relationship between k and model accuracy. To address this, we tested multiple k values and selected the best one, as performed in the DNABERT study. Building on this, we now integrate multi-scale k-mer statistics, spanning from k=3 to k=6. We use both RF and CNN models in this approach.

For the RF model, we maintain the same tabular format as before, but this time we incorporate multiple k-mers into a unified feature set. In the case of the CNN model, an ensemble approach was used. Each individual CNN processes a separate k-mer size (k=3, k=4, k=5, and k=6) with the same architecture as previously used, but with the last layer removed. Instead of the original output layer, the outputs from each model are flattened and passed through a fully connected layer (metamodel), and then through a final sigmoid neuron to generate the final prediction. The dimension of the fully-connected layer was optimized separately for each species using grid search, using values ranging from 32 to 512. This multiscale integration approach, while non-trivial for DNABERT, is straightforward in our framework and effectively combines the complementary strengths of different k-mer sizes.

However, feature importance analysis reveals that when specific k-mers are significant, their related variants (e.g., subsets or extensions) may also appear important, indicating potential multicollinearity. While this could affect some classifiers, it is not a major issue for RF and CNN in practice.

Our results, summarized in Table VI, represent the average performance from stratified 5-fold cross-validation. The multi-k RF and CNN models consistently match or exceed DNABERT’s performance, while being far less computationally demanding. In addition, our approaches demonstrate a notable advantage on smaller datasets, such as those for *A. thaliana, G. max* and *O. sativa*, as well as on datasets with longer sequences, including *B. rapa* and *G. max*. These findings are particularly encouraging given the challenges associated with large-scale genomic studies. Access to extensive datasets is often limited, and lncRNAs are known for their considerable length, which poses a well-documented challenge for LLM-based models like DNABERT, due to their context length limitations.

**TABLE VI.**
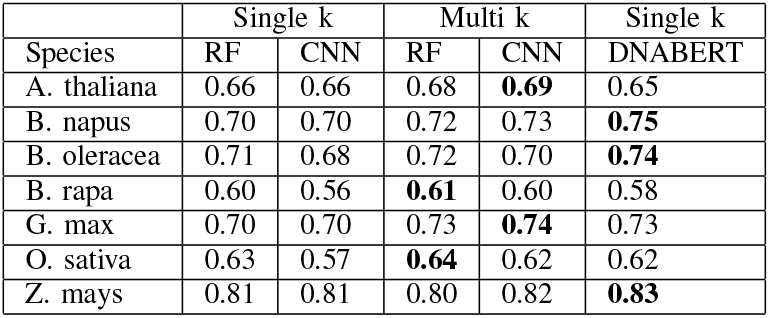
Comparison of classification accuracy for single-k-mer models, multi-k-mer models(with*k* ranging from3 to6), and the best-performingdnabert model across seven plant species. the results represent the mean performance obtained from stratified5-fold cross-validation.

## V. Conclusion and future work

This study presents a novel approach for extending FCGR with positional information (PFCGR) by incorporating statistical measures of k-mer positions, such as mean, standard deviation, skewness, and kurtosis. In this way, our method improves the standard FCGR by adding layers of positional information, providing a richer set of features for the classifiers. We combine this approach with machine-learning techniques, such as Logistic Regression (LR), Random Forest (RF) and Convolutional Neural Networks (CNN), to classify lncRNAs of seven crop species. Our method reduces computational time by 80% to 95% compared to the state-of-the-art DNABERT for this task, while maintaining competitive accuracy, and even in some instances surpassing it.

In addition, our experiments across multiple crop species illustrate the robustness and adaptability of our approach. Notably, the RF and the CNN have shown significant promise, leading to high accuracy and improved classification performance, particularly when using the multi-k feature sets. However, we note that applying our approach to other types of sequences and integrating more diverse datasets are necessary to validate and potentially improve the generalizability of our findings. Importantly, the PFCGR methodology is sequence-agnostic and can be readily extended to any DNA, RNA or protein, assuming the protein-adapted FCGR is used, sequence classification task, making it a versatile tool for diverse applications.

Lastly, we want to underline that this study not only advances the computational tools available for genomic research, but also contributes to our understanding of lncRNA functions and their regulatory roles in plants. By providing a more efficient and accurate tool for lncRNA identification, we pave the way for discoveries in plant biology, improve prospects for agricultural innovation, and accelerate plant breeding programs.

In future work, we can focus on exploring alternative neural network architectures, such as transformers, which could improve accuracy, especially with larger datasets. Additionally, incorporating gapped k-mers could improve generalizability and robustness by accommodating sequencing errors and variations. These strategies will further improve the accuracy and versatility of our method.

## VI. Acknowledgments

This work was supported by the National Institutes of Health (NIH) under grant number 5R01HG011795 and by the Cancer Prevention and Research Institute of Texas (CPRIT) under grant number RP240131.

